# Neutrophil Reprogramming Underlie Vasculopathy and Lung Disease in Systemic Sclerosis

**DOI:** 10.1101/2025.02.12.637866

**Authors:** Norma Maugeri, Giuseppe A. Ramirez, Annalisa Capobianco, Antonella Monno, Marco E. Bianchi, Patrizia Rovere-Querini, Marco Matucci-Cerinic, Angelo A. Manfredi

## Abstract

The role of neutrophils in systemic sclerosis (SSc) remains incompletely understood. To address this, blood samples from 39 SSc patients, 39 healthy controls, and 22 systemic lupus erythematosus (SLE) patients were analyzed. In SSc, neutrophils exhibited substantial activation, evidenced by granule mobilization, elevated plasma levels of Neutrophil Extracellular Trap (NET) byproducts, and upregulated TIE2 expression. In parallel, they underwent metabolic reprogramming, characterized by increased autophagy, likely to support the heightened energy demands of activation. By contrast, neutrophils from SLE patients displayed minimal autophagy, lacked TIE2 expression, and shifted toward low-density granulocytes. Neutrophil reprogramming in SSc correlated with plasma levels of HMGB1^+^ EVs. Mechanistically, EVs purified from the plasma of patients with SSc adhered to neutrophils when injected in immunodeficient NSG mice, inducing autophagy, TIE2 expression, and promoting lung inflammation and fibrosis. These effects were abrogated by HMGB1 inhibitors and required the HMGB1 receptor, RAGE. Recombinant HMGB1 recapitulated EV-induced effects, while neutrophil targeting by liposome-encapsulated clodronate prevented them. In summary, neutrophils in SSc exhibit a dual phenotype of autophagy and activation driven by HMGB1^+^ EVs, representing a pathogenic mechanism and a potential therapeutic target in SSc.

**Graphical abstract:** 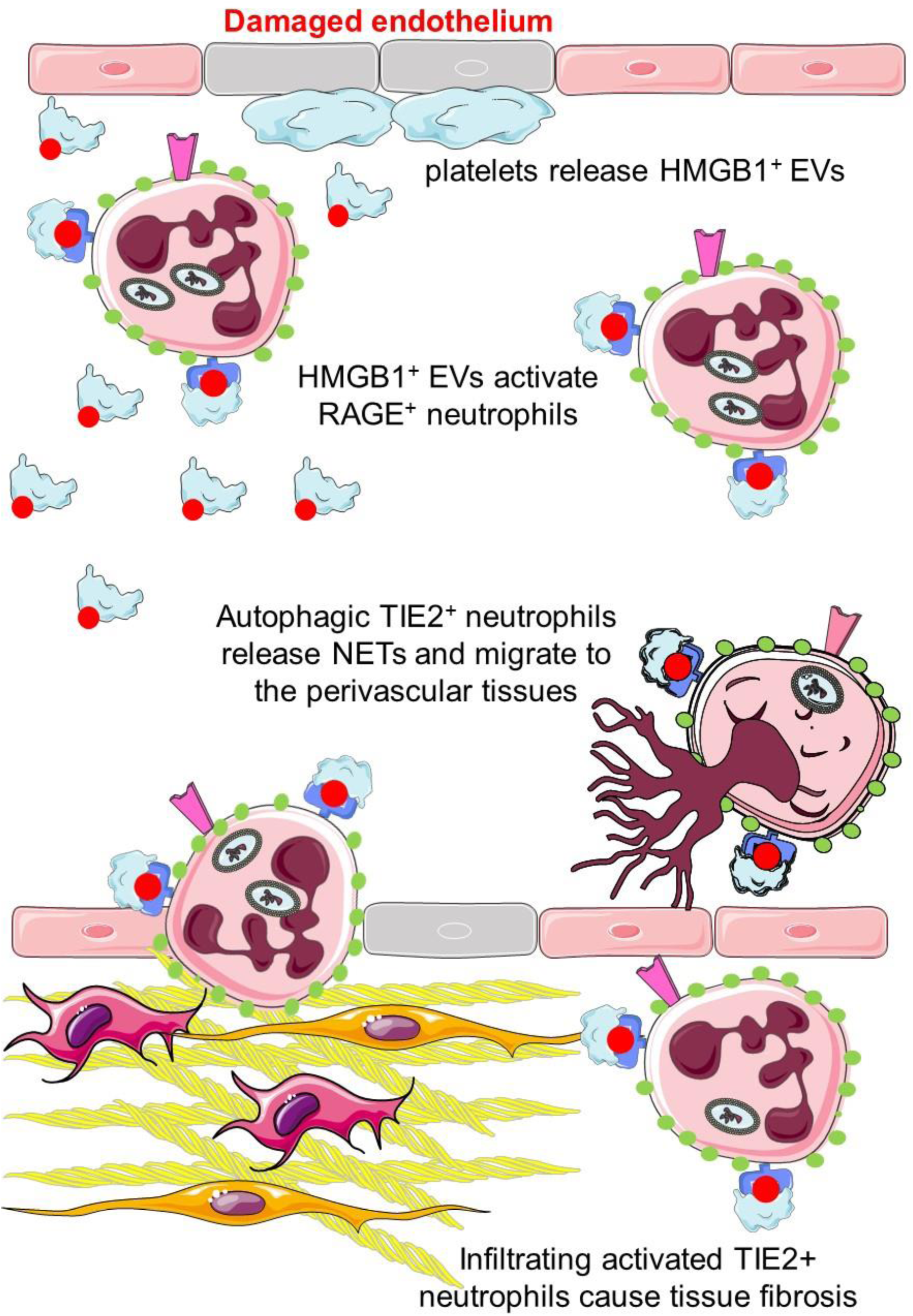

## Introduction

Systemic sclerosis (SSc) is characterized by widespread vascular abnormalities and progressive fibrosis of the skin and internal organs. Therapeutic options remain limited, underscoring the need for translational research to bridge the gap between basic science discoveries and clinical applications (1, 2). Early events in SSc natural history include endothelial cell activation and death, capillary loss, and exposure of sub-endothelial tissues. This results in progressive thickening of the media and hyperplasia of the intima, eventually leading to lumen obliteration (2).

Platelets activation and degranulation has been described across patient cohorts (3). It contributes to microvascular damage and vessel wall remodeling (4–7) and is closely associated with neutrophil activation, which significantly influences gene expression in patients’ blood samples (3, 8, 9). Normalization of neutrophils gene expression correlates with the improvement in lung function observed following hematopoietic stem cell transplantation (10). Functional studies have also directly implicated neutrophils activation and functional exhaustion in the natural history of SSc (11, 12). In two independent prospective SSc cohorts, blood neutrophil count and the neutrophil-to-lymphocyte ratio, which do not account for the heterogeneity of neutrophil composition, predicted an increased mortality and more severe disease (10). Byproducts of NETs, that reflect neutrophil activation, accumulate in SSc plasma (7, 13–15) and are correlated with biomarkers of vascular injury, including soluble E-selectin and VCAM-1 (16), and with endogenous neutrophil agonists, such as mitochondrial-derived N-formyl methionine (17).

The mechanisms underlying neutrophil involvement and its association with vasculopathy remain elusive in SSc. In contrast, significant advancements over the past years have elucidated the multifaceted roles of neutrophils in other systemic autoimmune diseases. Notably, in systemic lupus erythematosus (SLE), neutrophils are well known to contribute to vascular dysfunction and homeostasis disruption, while fostering the autoimmune response by releasing oxidized nucleic acids and post-translationally modified histones into the extracellular space (18). Furthermore, extracellular nucleic acids in SLE amplify type I interferon production by plasmacytoid dendritic cells, perpetuating the inflammatory cycle (18).

Extracellular vesicles (EVs) are membrane-bound structures released during cell activation or stress (7, 14, 19). EVs accumulate in the blood of patients with SSc and are biologically active, contributing to key features of the disease such as lung fibrosis (15). At least in part their bioactivity depends on the ability to present HMGB1, a potent agonist, to neutrophils and vascular cells (7, 14, 20).

In this study, we show that EVs accumulating in the blood of SSc patients trigger neutrophil reprogramming, inducing autophagy, upregulation of the angiopoietin receptor TIE2, and the formation of NETs in a redox state-dependent manner. Furthermore, SSc EVs expressing HMGB1 induce in vivo lung fibrosis which can be prevented by depleting TIE2-expressing neutrophils. This may therefore emerge as a new potential treat to target pathway in SSc.

## RESULTS

### Autophagy and activation of neutrophils of patients with SSc

We compared neutrophils in the blood of patients with SSc to those of healthy volunteers. As a reference group, we studied patients with SLE, a disease in which dysregulated neutrophil activation plays a role in vascular disease (21). However, unlike SSc, fibrosis and obliterative vasculopathy are uncommon in SLE. **Tables 1** and **2** report the main characteristics of the patients.

Most blood neutrophils in SSc patients, but not in healthy donors, were activated, as indicated by the localization of MPO to the plasma membrane, a *bona fide* evidence of azurophilic granules mobilization (**Figure 1**, panels **A** and **B**). Additionally, most neutrophils were autophagic, as assessed by the accumulation of the Cyto-ID tracer within autophagosomes (**Figure 1**, panels **A** and **C**), a feature linked to prolonged survival and extended biological action.

**Figure 1:**
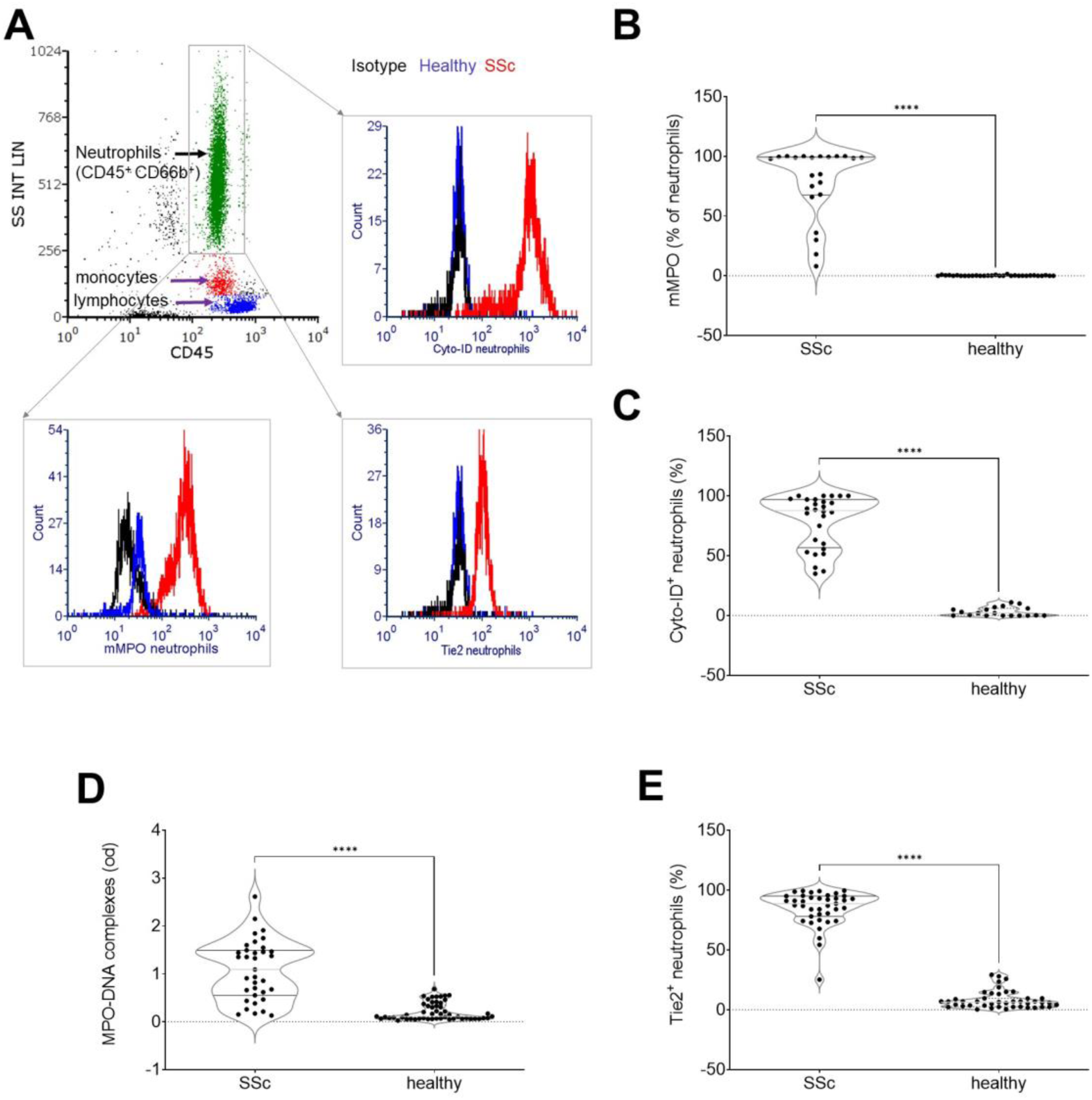
Neutrophil Activation in Systemic Sclerosis. A. Neutrophils were identified among blood leukocytes by flow cytometry, using side scatter (SS), and the expression of the pan-leukocyte marker CD45 alongside the neutrophil­specific marker CD66b. In representative SSc patients (red profiles), neutrophils exhibited significant autophagy, indicated by Cyto-ID dye accumulation in autophagosomes, displayed surface expression of MPO, and upregulated the angiopoietin receptor, TIE2. Results for neutrophils from representative healthy controls (blue profiles) and with isotype-matched control antibodies (black profiles) are also shown. Comparison between SSc patients and healthy controls reveals significantly increased surface expression of MPO (B), elevated autophagy (assessed by Cyto-ID dye accumulation, C), enhanced NET generation (quantified by MPO-low molecular weight DNA complexes, D), and heightened expression of the activation marker T1E2 in neutrophils from SSc patients (E). ****, significant different from healthy subjects, p < 0.001.

Autophagy also provides the necessary energy to produce NETs (22–24). Indeed, complexes of low molecular weight DNA fragments with MPO, a marker for NET formation, were significantly more concentrated in the plasma of SSc patients (**Figure 1**, panel **D**). Among the markers indicating neutrophil activation, we assessed the expression of the angiopoietin receptor, TIE2, that has been involved in neutrophil activation and production of NETs (25, 26).

In SSc patients, a significant proportion of neutrophils exhibited elevated TIE2 expression on their plasma membranes (84.7 ± 2.4%) compared to healthy donors (8.3 ± 1.1%; p < 0.0001) (**Figure 1**, panel **E**). Immunogold electron microscopy further confirmed TIE2 localization within extra-granular cytoplasmic domains in SSc neutrophils, contrasting with its prevalent localization on the plasma membrane in neutrophils from healthy donors (**Supplementary Figure,** compare panel **F1** with panels **F2-F4)**.

Neutrophils from SLE patients also exhibited enhanced MPO membrane expression while NETs fragments accumulated in the patients’ plasma (**Figure 2**, panels **A**-**B**). However, neutrophils in SLE and SSc patients were markedly different: the extent of neutrophil autophagy and the proportion of TIE2-expressing neutrophils were significantly lower in SLE patients (**Figure 2**, panels **C**-**D**). As previously reported (27, 28), low-density neutrophils (LDNs) were a prominent subset within the peripheral blood mononuclear cell fraction isolated from systemic lupus erythematosus (SLE) patients using density gradient centrifugation. In stark contrast, LDNs constituted only a minority of the mononuclear cells in SSc patients (**Figure 2**, panel **E**). Notably, the LDNs from SSc patients were autophagic, a feature absent in LDNs from SLE patients.

**Figure 2:**
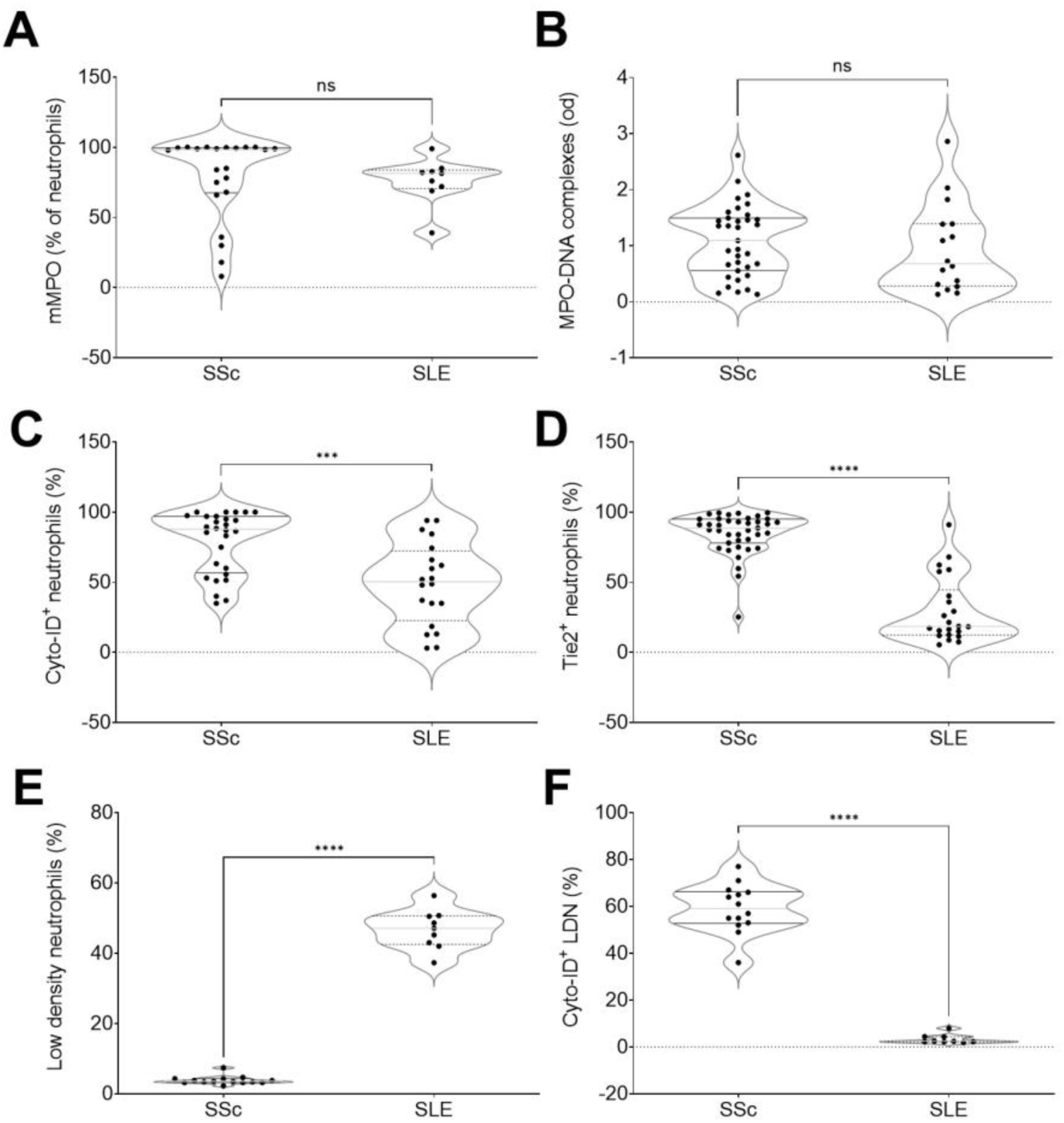
Different features of neutrophils activation in SLE and SSc. Comparison of neutrophil activation between patients with SSc and SLE reveals similar levels of primary granule content mobilization, as indicated by membrane expression of MPO (mMPO, A) and comparable NET generation, quantified by MPO-low molecular weight DNA complexes **(B).** However, SSc patients exhibited significantly higher autophagy (assessed by Cyto-ID dye accumulation, **C)** and greater expression of the activation marker, TIE2 (D). Additionally, Low-Density Neutrophils (LDN) were less abundant in SSc patients compared to SLE (E). Notably, in SSc LDNs were autophagic, whereas in SLE these cells showed no signs of autophagy (F). ***‘, significant different from healthy subjects, p < 0.001; "*, significant different from healthy subjects, p < 0.005. ns: non-significant.

Given the neutrophil short half-life, it is likely that the stimulus driving their reprogramming persists in the SSc patient’s bloodstream. We have previously observed that HMGB1^+^ EVs trigger neutrophil activation, promote autophagy, and induce the production of NETs (7, 14, 24, 29). HMGB1^+^ platelet-derived EVs (plt-EVs), identified based on the expression of the platelet lineage marker CD61 (GPIIbIIIa), were more concentrated in the blood of SSc patients compared to controls (p < 0.0001, **Figure 3**, panel **A**), and their levels significantly and strongly correlated with the expression of the TIE2 receptor on activated neutrophils and with the extent of autophagy (**Figure 3**, panels **C-D**), supporting the possibility of a causal link.

**Figure 3:**
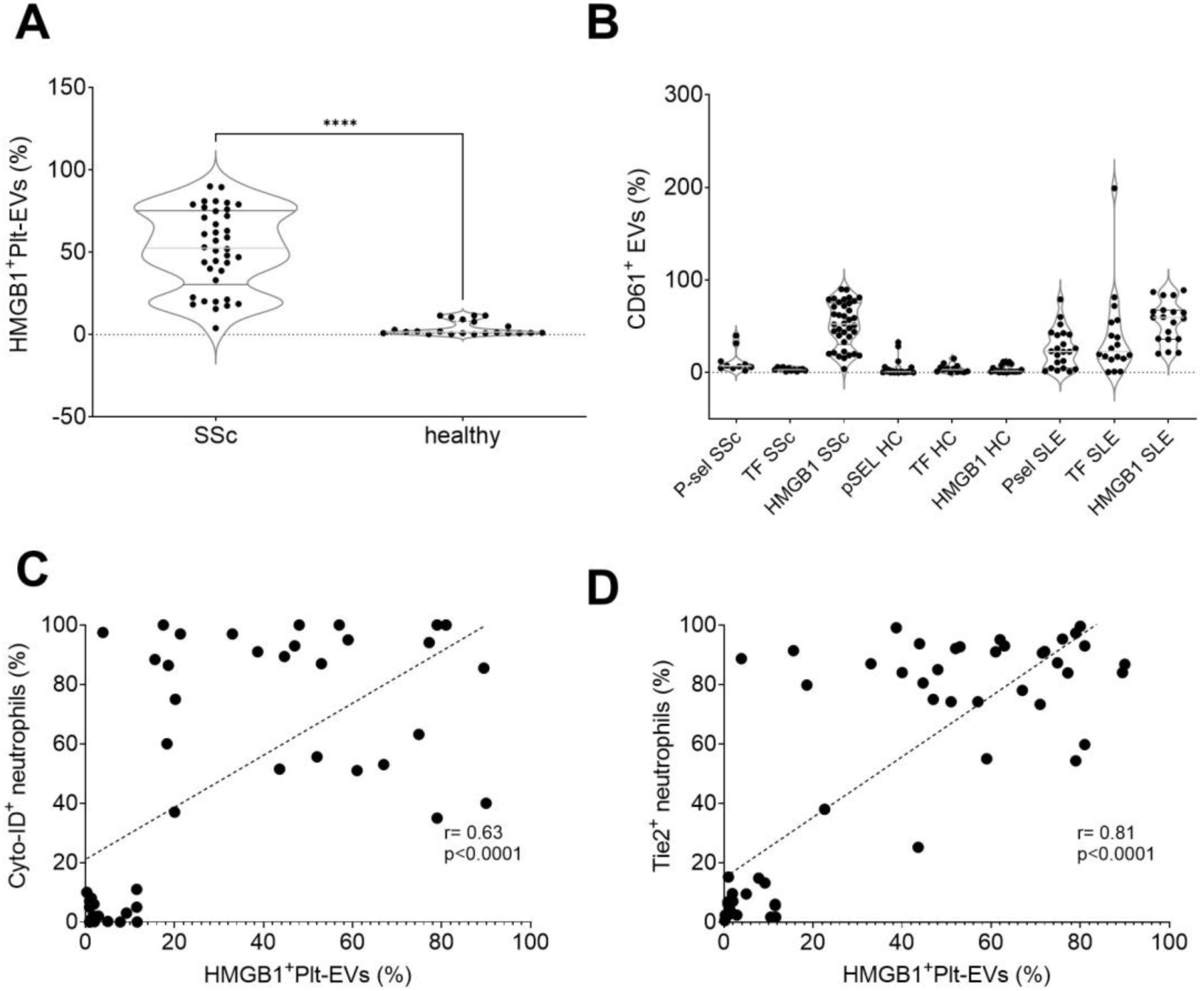
Accumulation of platelet HMGBV EVs correlates with neutrophil activation in SSc. Platelet-derived HMGB1’ EVs (HMGB1^1^ pit EVs) were significantly more represented in the blood of SSc patients compared to healthy controls (A). In contrast, there were no significant differences in platelet-derived EVs expressing P-selectin (Psel) or tissue factor (TF) **(B).** HMGB1* pit EVs (x-axis) showed a significant positive correlation with both neutrophil autophagy (C, y-axis) and TIE2 expression **(D,** y-axis). ****, p < 0.001.

In contrast, other signals potentially involved in EV interactions with the innate immune system, particularly with neutrophils—such as P-selectin—or in the activation of the coagulation system, like tissue factor were not significantly overexpressed on SSc EVs (**Figure 3**, panel **B**) and the concentration of P-selectin^+^ or TF^+^ EVs showed no correlation with the extent of neutrophil activation or autophagy.

### Vesicular HMGB1 causes neutrophil TIE2 upregulation

To elucidate the causal relationship between EVs and neutrophil activation in SSc, we exposed normal neutrophils to EVs isolated from the plasma of SSc patients. Given that platelets are a primary source of EVs in SSc patients (7, 14, 30), we also generated EVs in vitro by activating platelets. Healthy neutrophils challenged with SSc EVs reproduced most features observed in neutrophils of SSc patients, including MPO expression on the neutrophil plasma membrane, expression of the TIE2 receptor, and accumulation of Cyto-ID dye in autophagosomes (**Figure 4**, panels **A-C**). Activation of neutrophils could also be detected by assessing the generation of NETs by autophagic neutrophils, either by measuring the concentration of MPO-DNA complexes in the supernatant or by fluorescence microscopy (**Figure 4**, panels **D** and **E**).

**Figure 4:**
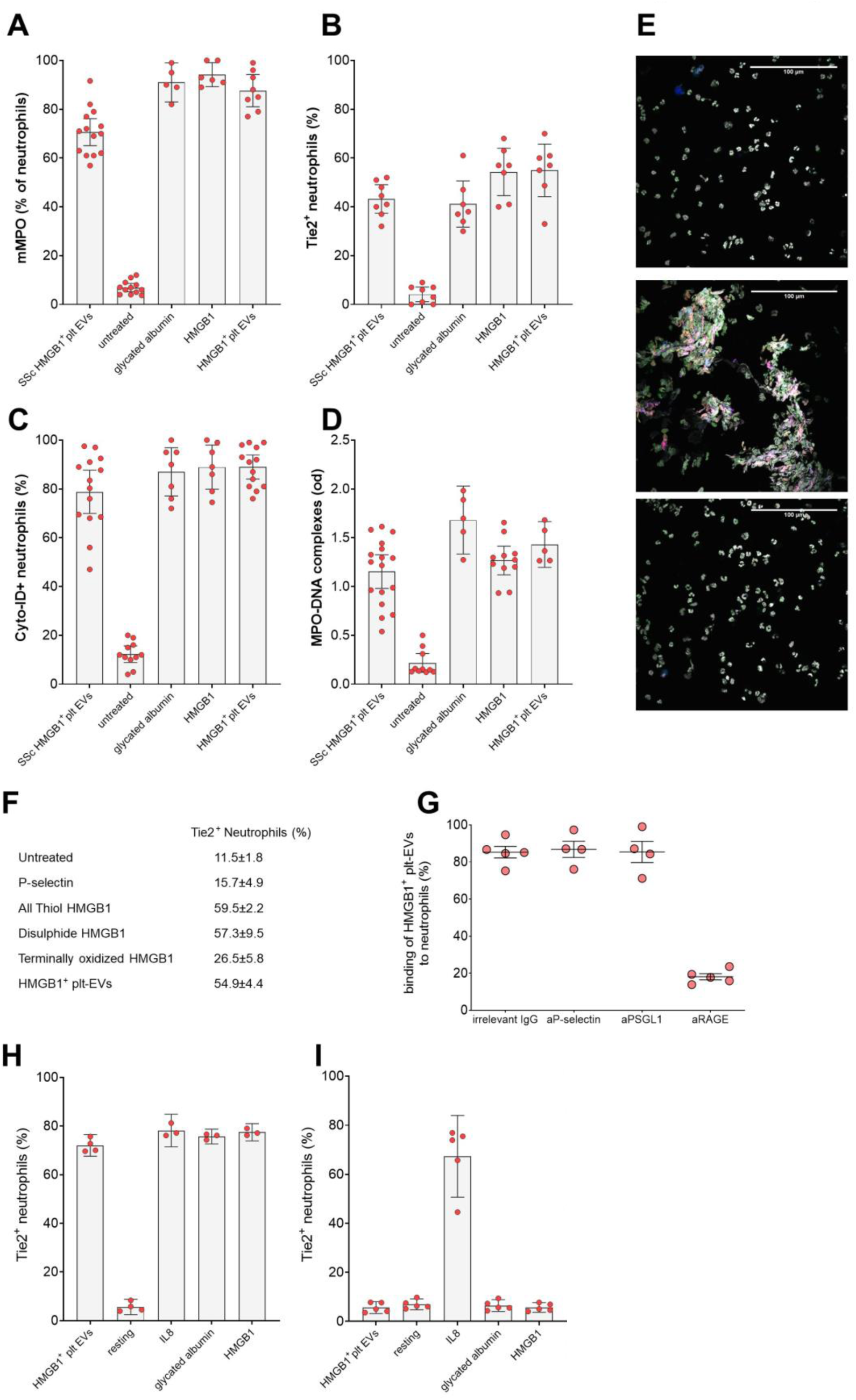
HMGBl^EVs induce neutrophil activation through a RAGE-dependent pathway-Purified human neutrophils were either challenged with platelet-derived IIMGB1* EVs from the plasma of SSc patients (SSc 1IMGB1^+^pit EVs), left untreated, or stimulated with the RAGE agonist glycated albumin, with recombinant HMGB1, or HMGB1* EVs from activated platelets (HMGBVplt EVs). Neutrophil activation was assessed by evaluating: membrane-bound MPO (mMPO, A), expression of the activation marker TIE2 (B), (extent of autophagy (measured via Cyto-ID dye accumulation, C), and NET generation (quantified by MPO-DNA complex levels in the supernatant D). E: Neutrophils were either left untreated (top panel) or exposed to platelet-derived HMGBI^+^ EVs in the absence (middle panel) or the presence (bottom panel) of a mAb blocking RAGE. Cells were then analyzed by confocal microscopy for autophagy (Cyto-ID dye, green), TIE2 (Alexa 647, blue), citrullinated H4 (Alexa 546, red). DNAwas counterstained with Hoechst (white). F: Neutrophils were either left untreated or exposed to purified human P-selectin, redox isoforms of HMGB1 (all thiol, disulphide, or terminally oxidized), or platelet-derived HMGB1+ EVs. The expression of the T1E2 receptor on neutrophils was assessed by flow cytometry. G: Neutrophils were treated with isotype-matched control antibodies or antibodies specific for P-selectin (aPselectin), its counter-receptor on^1^ neutrophils PSGL1 (aPSGLI), or the HMGB1 receptor RAGE (aRAGE) The interaction between platelet-derived HMGBT EVs and neutrophils (y-axis) was measured by flow cytometry. Neutrophils from wild-type (H) and RAGE­deficient (I) mice were exposed to platelet-derived HMGBl^+^EVs, left untreated, or challenged with the neutrophil agonist IL-8, the RAGE agonist glycated albumin, or recombinant HMGB1. TIE2 receptor expression on neutrophils was assessed by flow cytometry.

Moreover, purified recombinant human HMGB1 reproduced the effects of the EVs, as did glycated albumin, a known agonist of the HMGB1 receptor RAGE (**Figure 4**, panel **A**), suggesting that the HMGB1-RAGE interaction may be involved. Indeed, interference with RAGE effectively abrogated the interaction of HMGB1^+^ plt-EVs with neutrophils, while blocking P-selectin or the PSGL1 counter-receptor on neutrophils did not affect plt-EV adhesion **(Figure 4**, panel **F)**. The action of recombinant HMGB1 in triggering neutrophil TIE2 upregulation was dependent on the molecule’s redox state (**Figure 4**, panel **G**). Notably, purified P-selectin alone was ineffective at modulating TIE2 expression in neutrophils (**Figure 4**, panel **G**).

Additionally, HMGB1^+^ plt-EVs did not induce TIE2 expression in mouse neutrophils lacking the HMGB1 receptor, RAGE (**Figure 4**, panels **H** and **I**). Neutrophils from *Rage*⁻/⁻ mice upregulated TIE2 in response to the RAGE-independent agonist IL-8, demonstrating their ability to undergo activation. However, unlike wild-type neutrophils, *Rage⁻/⁻*neutrophils failed to upregulate TIE2 when exposed to recombinant HMGB1 or the other RAGE agonist, glycated albumin (**Figure 4**, panels **H** and **I**). This indicates that RAGE is necessary for mediating plt-EVs HMGB1’s effects on neutrophils.

EVs purified from the plasma of SSc patients were injected into the tail vein of immunodeficient NSG mice, which are receptive to grafting of human cells and tissues. SSc-derived, CD61^+^ plt-EVs adhered to host circulating neutrophils, with a substantial proportion (84.5 ± 5.3 %) of EV-neutrophil aggregates detectable in the blood six hours after injection (**Figure 5**, panel **A**). The interaction persisted over time, with most circulating neutrophils remaining associated with EVs eighteen hours after injection. Importantly, neutrophils interacting with SSc EVs selectively upregulated the expression of TIE2, and this event was abrogated by HMGB1 inhibitors, Box A and low molecular weight heparin (LMWH; **Figure 5**, panel **A**). Adhesion to neutrophils was negligible when we injected into NSG mice EVs purified from healthy subjects, that expressed very limited amounts of HMGB1 (**Figure 3**, panel **A**). Furthermore, neutrophils of mice injected with EVs from healthy donors failed to upregulate TIE2 expression following the injection of EVs (**Figure 5**, panels **A-B**).

**Figure 5:**
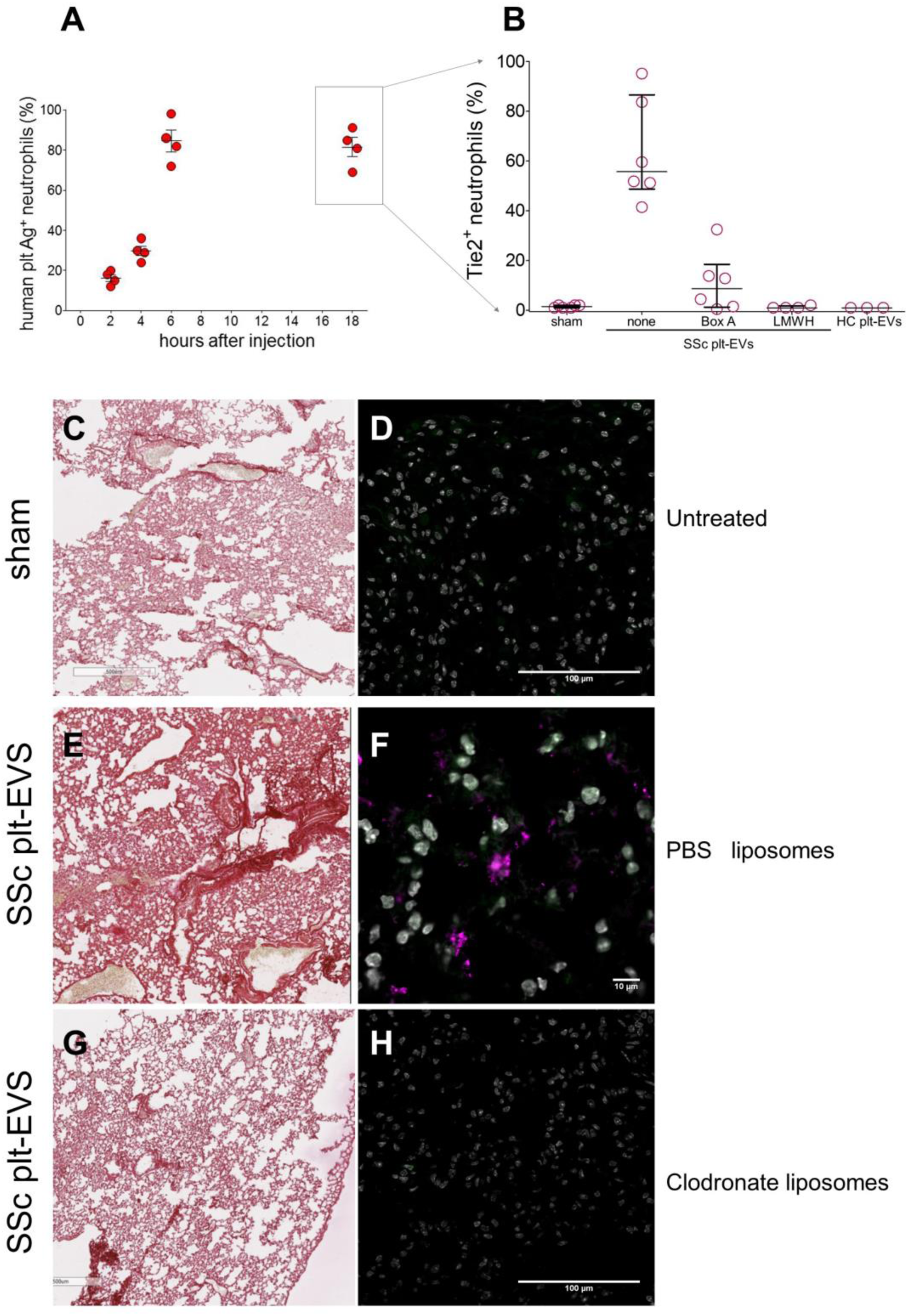
Neutrophils mediate SSc-HMGBl∼ EVs-induced inflammation and fibrosis. EVs from SSc patients and controls were injected into the tail vein of NSC mice. A. The binding of human platelet-derived EVs to neutrophils (% on y-axis) was assessed over time (hours, x-axis) post­injection. B. Neutrophil TIE2 expression (y-axis) was measured 18 hours post-injection in control (sham) mice, in mice injected with SSc platelet-derived EVs (SSc plt-EVs), and in mice receiving SSc plt-EVs alongside HMGB1 inhibitors (BoxA or LMWH). A control group of mice was injected with EVs from healthy subject plasma (HC plt-EVs). C-H. Lungs were collected post-euthanasia from sham-treated control mice (C, D) and mice injected rath SSc plt-EVs. Comparisons were made between mice pre-treated with saline liposomes (PBS liposomes, E-F) and those pre-treated with clodronate liposomes to inhibit neutrophils (G-H). Panels C, E, and G show histological analysis with hematoxylin and eosin staining, while panels D, F, and H depict confocal microscopy findings. Neutrophils (green), TIE2 (Alexa 647, pink), and DNA was counterstained with Hoechst (white).

EVs purified from the blood of SLE patients injected in immunodeficient NSG mice demonstrated a similar limited efficacy in modulating blood neutrophil characteristics and causing endothelial damage. Specifically, they failed to induce significant changes in neutrophil autophagy, granule mobilization, NET formation, or the concentration of soluble E-selectin compared to levels observed in saline-treated mice (**Table 3**).

**Table 1:**
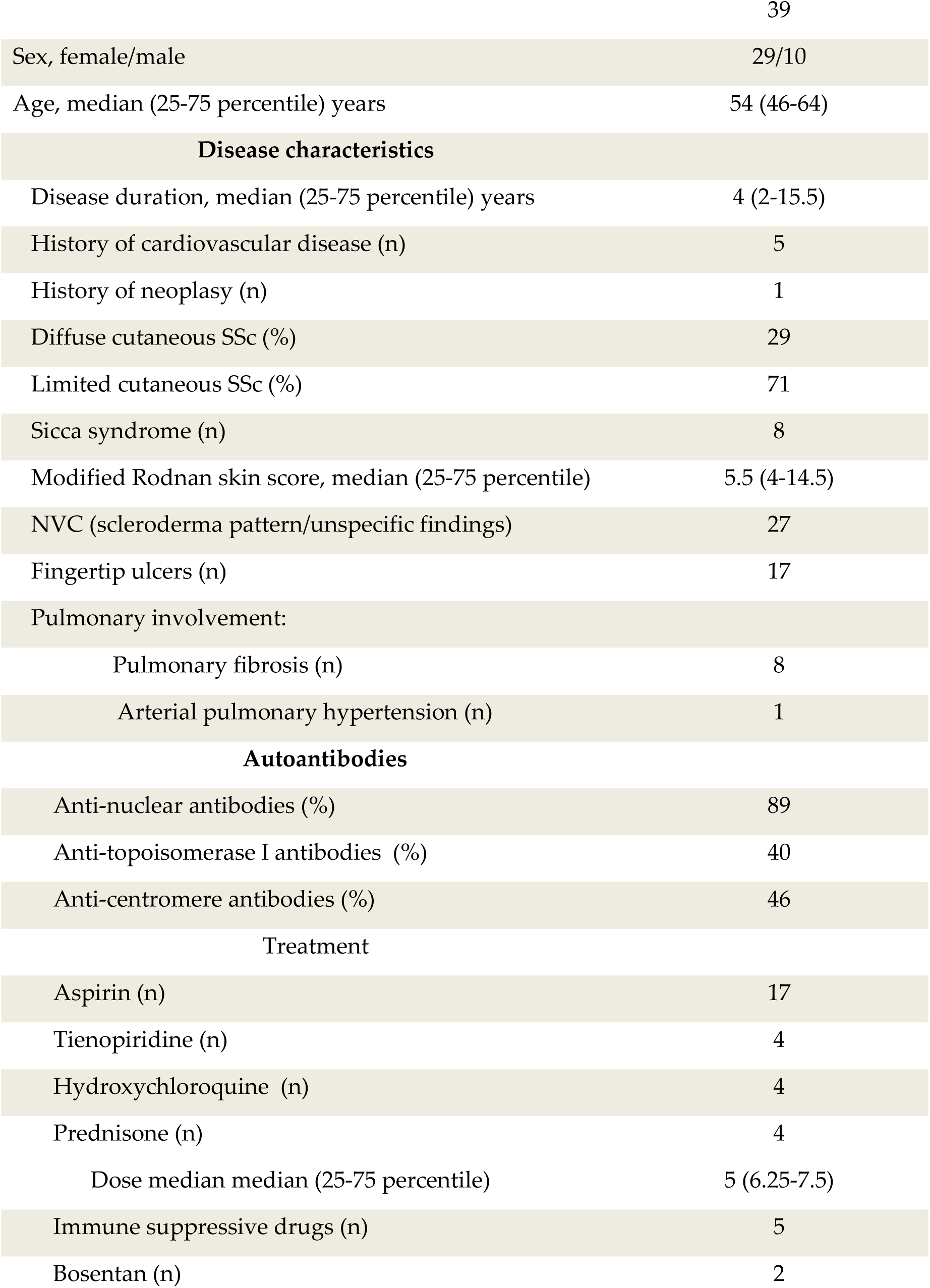
SSc patients’ characteristics.

|  |  |
| --- | --- |
|  | 39 |
| Sex, female/male | 29/10 |
| Age, median (25-75 percentile) years | 54 (46-64) |
| **Disease characteristics** |  |
| Disease duration, median (25-75 percentile) years | 4 (2-15.5) |
| History of cardiovascular disease (n) | 5 |
| History of neoplasia (n) | 1 |
| Diffuse cutaneous SSc (%) | 29 |
| Limited cutaneous SSc (%) | 71 |
| Sicca syndrome (n) | 8 |
| Modified Rodnan skin score, median (25-75 percentile) | 5.5 (4-14.5) |
| NVC (scleroderma pattern/unspecific findings) | 27 |
| Fingertip ulcers (n) | 17 |
| Pulmonary involvement: |  |
| Pulmonary fibrosis (n) | 8 |
| Arterial pulmonary hypertension (n) | 1 |
| **Autoantibodies** |  |
| Anti-nuclear antibodies (%) | 89 |
| Anti-topoisomerase I antibodies (%) | 40 |
| Anti-centromere antibodies (%) | 46 |
| Treatment |  |
| Aspirin (n) | 17 |
| Tienopiridine (n) | 4 |
| Hydroxychloroquine (n) | 4 |
| Prednisone (n) | 4 |
| Dose median median (25-75 percentile) | 5 (6.25-7.5) |
| Immune suppressive drugs (n) | 5 |
| Bosentan (n) | 2 |

**Table 2:** SLE patients’ characteristics.

|  |  |
| --- | --- |
| N | 22 |
| Sex, female/total | 17/22 |
| Age, median (25-75 percentile) years | 42 (33-52.3) |
| **Disease characteristics** |  |
| Disease duration, median (25-75 percentile) years | 14 (6-22.5) |
| SLEDAI, median (25-75 percentile) | 2 (2-6) |
| Renal involvement (n) | 11 |
| Haematological involvement (n) | 12 |
| Neurological involvement | 3 |
| Antiphospholipid syndrome (n) | 1 |
| **Autoantibodies** |  |
| Anti - ds DNA | 19 |
| Anti - Sm | 5 |
| Anti- phospholipid | 10 |
| **Treatment** |  |
| Aspirin (n) | 5 |
| Tienopiridine (n) | 1 |
| Warfarin (n) | 2 |
| Hydroxychloroquine (n) | 11 |
| Prednisone (n) | 5 |
| Dose: median (25-75 percentile) | 5 (2.5-5) |
| Methotrexate (n) | 3 |
| Mycophenolate mofetil | 4 |

**Table 3:** HMGB1-Positive Extracellular Vesicles from SSc Patients cause the In Vivo Activation of Neutrophils and Endothelial Cells.

|  | Autophagy<br>(Cyto-ID+,<br>% of<br>neutrophils) | Granules<br>mobilization<br>(mMPO,<br>% of<br>neutrophils) | NETs<br>(plasma<br>citH4, OD) | Endothelial<br>activation<br>(plasma sE-<br>selectin,<br>ng/mL) |
| --- | --- | --- | --- | --- |
| Sham-treated | 16±5 | 11±1 | 0.07±0.03 | 0.8±0.06 |
| SSc-EV | 62±4* | 78±2* | 0.49±0.05* | 224±43.5* |
| SSc-EV + HMGB1 blockade | 11±3 | 14±2 | 0.07±0.01 | 4.5±1.2 |
| SLE-EV | 27±6 | 18±6 | 0.07±0.02 | 2.0±1.2 |
| SLE-EV + HMGB1 blockade | 38±13 | 19±3 | 0.09±0.02 | 1.4±0.6 |

\*: $p < 0.001$ , significantly different from mice treated with saline (sham-treated), from mice treated with SSc-EV in the presence of HMGB1 blockade with the competitive antagonist, BoxA, and from mice treated with SLE-EV.

After the injection of SSc-EVs, TIE2-expressing neutrophils infiltrated the lung parenchyma, resulting in alveolar obliteration and diffuse fibrotic remodeling associated with tissue injury. In contrast lung sections from the saline-treated (sham-treated) group displayed no evidence of ongoing inflammatory responses, alveolar damage, or interstitial fibrosis (**Figure 5**, panels **C**-**F**).

Selective neutrophil stunning using liposome-encapsulated clodronate, which interferes with early neutrophil migration, cytokine and chemokine production, and generation of NETs (31) prevented inflammatory changes and subsequent lung damage and remodeling in mice injected with SSc-EVs. Importantly, the lung disease elicited by SSc-EVs was unaffected in mice injected with liposomes containing saline, confirming that the effect of EVs in vivo was mediated via their action on neutrophils (**Figure 5**, panels **E**-**H**).

## Discussion

In this study, we demonstrate that neutrophils in patients with SSc exhibit distinct features of aberrant activation and autophagy, which may be integral to the disease’s progression. Neutrophils play a role in responding to vascular injury by aiding in endothelial repair and clearing harmful agents. Under normal conditions, their activity subsides once the causative factor is resolved and the tissue has healed. However, when initial damage fails to resolve, persistent and excessive activation may occur, perpetuating vessel damage through sustained generation of inflammatory molecules and the destructive effects of NETs (17).

In SSc, a vicious cycle of inflammation, tissue injury, and NET formation may stem from the inability of blood vessels to properly repair themselves (32). This impaired repair likely involves dysregulated angiopoietin/TIE2 receptor signaling between activated endothelial cells, perivascular cells, and potentially inflammatory leukocytes. Vasculopathy in SSc occurs independently of the extent of fibrosis and significantly contributes to morbidity (17). Neutrophils are likely involved in the initiation and maintenance of the persistent and widespread microvascular involvement observed in SSc vasculopathy, as suggested by findings in other conditions (17, 33, 34). However, despite mounting recent evidence involving neutrophils in SSc pathophysiology (11, 16, 35–38), there is still limited detailed data regarding their actual roles. Our findings provide new insight into this gap, demonstrating a widespread reprogramming of neutrophils in consecutive patients with SSc.

The reprogramming observed in neutrophils from SSc patients prompted us to investigate the stimuli driving this phenomenon. EVs were a promising candidate, since they could interact with neutrophils in the circulation, influencing their interaction with the microvasculature. EVs accumulate in the blood of patients with SSc (7, 13, 15, 20, 30, 39–41), possibly because of defective phagocytic clearance of P-selectin expressing substrates (14). Most of EVs in SSc derive from platelets (13, 42) and are biologically active (7, 15, 20, 43–47). The connection between platelet activation, EVs accumulation, vascular injury and self-perpetuating vascular damage remains elusive.

We observe that a pivotal feature associated to the biological activity of EVs is the fact that they contain and present to the cells they interact with HMGB1, a potent mediator involved in vascular inflammation and tissue fibrosis (48, 49). HMGB1 has been previously shown to mediate inflammatory events, fibrosis included, through its receptor RAGE in a manner which is sensitive to the environmental redox balance (50), and we hypothesized that HMGB1^+^ EVs may be responsible for neutrophil activation in SSc.

Indeed, purified HMGB1^+^ EVs induced hallmark features of neutrophil metabolic and functional reprogramming. These included granule content redistribution, autophagy, NETs formation and TIE2 expression, all of which recapitulated the features observed in SSc patient neutrophils. Recombinant HMGB1 and the RAGE agonist, glycated albumin similarly induced neutrophil activation, suggesting that the HMGB1-RAGE axis may be a critical driver of this process. EVs purified from the plasma of SSc adhered to circulating neutrophils, with a persistent association which resulted in the selective upregulation of TIE2 on neutrophils. Blocking HMGB1 with inhibitors abrogated this response, confirming the central role of HMGB1 as a bioactive moiety presented by patients EV and responsible in driving TIE2 expression. Additionally, neutrophils from *Rage⁻/⁻* mice, which lack the receptor for HMGB1, did not upregulate TIE2 in response to HMGB1+ EVs, whereas they remained responsive to the RAGE-independent agonist, IL-8. This further underscores the critical role of RAGE in mediating HMGB1’s effects on neutrophils.

Notably, injection of SSc-derived EVs into NSG mice led to TIE2-expressing neutrophils infiltrating the lung parenchyma, where they caused significant alveolar obliteration and fibrotic remodeling. By “stunning” neutrophils with clodronate-containing liposomes (31) we prevented these events, confirming the importance of neutrophil infiltration in mediating lung damage.

Overall, our findings suggest that HMGB1^+^ EVs play a central role in driving neutrophil autophagy and activation in SSc. The upregulation of TIE2 on neutrophils and the subsequent infiltration into tissues contribute to the fibrotic and vascular manifestations of the disease. This study provides new insights into the mechanisms of neutrophil activation in SSc and identifies potential therapeutic targets to mitigate neutrophil-mediated tissue damage. Further research is needed to explore the therapeutic potential of targeting HMGB1 or TIE2 pathways in SSc. Of importance a role for RAGE in SSc associated vasculopathy has already been suggested, and high levels of the soluble molecule in patients with SSc at baseline may be used to predict new onset of pulmonary arterial hypertension and to predict lower survival (51). Understanding the precise mechanisms by which platelet-derived, HMGB1^+^ EVs interact with neutrophils and other immune and mural cells in SSc could open avenues for the development of novel targeted therapies.

### Patients and Methods

#### Patients

The study group consisted of thirty-nine patients classified according to the 2013 ACR/EULAR criteria for SSc (29 females) (52) (see **Table 1** for demographic and clinical characteristics). Thirty-nine age-matched healthy donors (median 51 years old, range 29-81; 21 females) served as controls. Twenty-two patients with SLE (17 females) classified according to the EULAR 2019 criteria (53) were studied in parallel. Patients with other systemic autoimmune disorders or overlap syndromes were excluded. All subjects gave their written informed consent to participate to the study. The Institutional Review Board approved the study.

#### Reagents

Monoclonal antibodies (mAbs) against CD45 (clone J33), CD61 (clone SZ21), CD66b (clone 80H3), CD62P (clone Thromb-6), PSGL-1 (clone PL-1), myeloperoxidase (MPO, clone CLB-MPO-1), relevant IgG isotype controls mAbs for flow cytometry, Thrombofix and Flow-Count™ Fluorospheres were obtained from Beckman Coulter (Italy). Monoclonal IgG1 isotype control (clone W3/25) was obtained from Acris Antibodies (Italy). mAb against HMGB1 (clone HAP-46.5), PGE1, Cell Death detection kit, glycated albumin, Thrombin Receptor Agonist Peptide-6 (TRAP-6) Sirius Red and Hoechst were obtained from Sigma (Italy).

Recombinant human interleukin (IL)-8 was from R&D (Italy). Zenon IgG Labeling kits (488, 546 or 647) were obtained from Invitrogen. Rabbit polyclonal anti-histone H4 (citrulline R3, ab81797) was obtained from Abcam (Prodotti Gianni, Italy). mAb anti human MPO used for the determination of DNA-MPO complexes was from ABD Serotec. BoxA and recombinant HMGB1 were obtained from HMGBiotech (Italy). The CYTO-ID® Autophagy detection kit was obtained by Enzo Life Sciences (3v Chimica, Milan, Italy). Monoclonal antibodies used to identify murine neutrophils (clone 7/4), and mAb anti RAGE (ab54741) antibodies were obtained from Abcam (Prodotti Gianni, Italy). Liposomes containing clodronate and liposomes containing PBS were obtained from ClodronateLiposomes.org (The Netherlands). mAb against TIE2 (clone 83715) for flow cytometry was from R&D (Italy) while the mAb against TIE2 (clone Ab33) for confocal and for electron microscopy was from Millipore (Italy).

#### Blood sampling

Venous blood was drawn through a 19-gauge butterfly needle. After having discarded the first 3-5 m, blood was carefully collected in tubes containing either EDTA to prepare PBMCs (54) (54), to purify EVs, platelets and neutrophils and to assess the concentration of NETs byproducts, or Na citrate and antiproteases for determination of cellular markers by flow cytometry (14, 29, 55).

#### Human blood cell preparations

Platelets, neutrophils and platelet EVs suspensions were prepared as described (7, 14, 20, 55–58). Briefly, blood samples were centrifuged at 150*g*, 10 min at 20°C, to obtain platelet-rich plasma to be used to isolate platelets while the remaining blood was used to purified neutrophils (see below). Briefly, the upper 2/3 of platelet rich plasma was recovered and PGE1 (2.5 µM) was added before centrifugation at 800*g* for 15 min at 18° C (without break). Platelet pellets were washed twice with Hepes Tyrode buffer (129 mmol/L NaCl, 9.9 mmol/L NaHCO3, 2.8 mmol/L KCl, 0.8 mmol/L KH2PO4, 5.6 mmol/L dextrose, 10 mmol/L HEPES, MgCl2 1 mM, pH 7.4) containing Na EDTA (5 mM) and in the first wash PGE1 (2.5 µM) and resuspended with Hepes Tyrode buffer containing CaCl_2_ (1 mM). Platelets were counted with an automated blood cell counter. Contaminating leukocytes were consistently absent. The purity of the platelet preparations was routinely confirmed by flow cytometry using anti-CD45 and anti-CD61 mAbs (<1 CD45^+^ event out of 10^7^ CD61^+^ events).

##### Extracellular vesicles

EVs were isolated from EDTA anticoagulated plasma samples and characterized as previously described (7, 14). Briefly, blood samples were centrifuged 15 min at 2,000 g at room temperature. Retrieved platelet-poor plasma was further centrifuged 15 min at 2,000 g and 4 °C to obtain platelet free plasma, then further centrifuged at 100,000g for 45 min at 4°C. EVs were retrieved in the pellets and resuspended in Hepes Tyrode buffer containing CaCl_2_ and MgCl_2_ (both 1mM final concentration). EV were also purified from the supernatant of purified platelets (5x10^5^/µL) stimulated with collagen (5µg/mL) for 60 minutes at 37°C in static conditions, then placed in chilled water bath for 2 min and centrifuged 15 min at 2,000 g and 4 °C. Supernatants were retrieved, centrifuged 45 min at 100,000xg at 4°C, pellets were resuspended in Hepes Tyrode buffer, and EVs expressing HMGB1 (HMGB1-EVs) were quantified by flow cytometry using flow count and diluted to a final concentration of 10^5^ HMGB1-EVs/µL.

##### Neutrophils

Neutrophils were isolated from the remaining blood (after the first centrifugation to obtain PRP) as previously described (59) by Dextran sedimentation followed by Ficoll-Hypaque gradient and hypotonic lysis of erythrocytes, were washed and resuspended in HEPES-Tyrode buffer with 1mM CaCl2. All procedures for neutrophil separation were performed at 4°C. The purified neutrophil suspensions were checked by flow cytometry to exclude the presence of monocytes and platelets (both virtually 0) as contaminants in the CD45+ CD66b+ populations.

##### Low density neutrophils

Frozen PBMCs suspensions were thawed, washed and fixed as previously described (54). After 4 hours at 4°C, samples were labelled with anti CD45 and anti CD66b and analyzed by flow cytometry to identify neutrophils within mononuclear cells.

#### Flow cytometry

All samples were analyzed on a diary aligned Navios flow cytometer (Beckman Coulter, Milan, Italy). Whole blood samples for the determination of cellular markers of activation were immediately fixed with equal volumes of Thrombofix, stored at 4°C and analyzed within 6-24 hours. Extent of platelet and leukocyte activation were assessed by multi parametric flow cytometry as previously described (20, 55, 58). Quantification of platelet-derived EVs (plt-EVs) in platelet-free plasma or in the supernatant of purified stimulated platelets was performed as described (7, 14, 57).

#### Neutrophil-EV interaction studies

Neutrophils freshly purified from the peripheral blood (5x10^3^/µL) were resuspended in Hepes Tyrode with CaCl_2_ and MgCl_2_ (both 1mM final concentration). Neutrophils were challenged at 37°C for 5 minutes with SSc-HMGB1^+^ plt-EVs, or with HMGB1^+^ plt-EVs, with recombinant HMGB1, with IL-8 or with glycated albumin. When indicated, plt-EVs were pretreated with blocking mAb against P-selectin, before the coincubation with neutrophils. Alternatively, neutrophils were pretreated with anti-PSGL-1 mAbs or with anti-RAGE before the incubation with platelets and plt-EVs. An irrelevant isotopic mAb was used as control. Reactions were stopped by addition of an equal volume of Thrombofix and samples stored at 4°C until analysis.

#### Mice

Ten-to twelve-week-old male NOD.Cg-Prkdcscid Il2rgtm1Wjl/SzJ mice (NSG) and *Rage*^-/-^ mice were obtained from Charles River (Milano). Our study examined male mice because male animals exhibited less variability in phenotype. Mouse experiments were performed in accordance with National and Institutional guidelines and experiments were approved by the Institutional Animal Care and Use Committee of the San Raffaele Scientific Institute (IACUC N°1216 and N°1371).

#### In vivo effects of EVs

EVs were injected in the tail vein of NSG mice (10 x 10^6^ EVs/mouse, as described (7, 14). Sham-treated mice were injected with Hepes Tyrode buffer (CaCl2 1mM). After 18 hs the blood was obtained from the orbital vein of the mice and the concentration of NETs byproducts (citrullinated Histone 4 and MPO-DNA complexes) and of plasma E-selectin assessed by ELISA (7, 14, 20, 29). Neutrophil stunning was obtained by treatments with liposomes containing clodronate (ClodronateLiposomes.org, The Netherlands) before EVs injection CC. Mice injected with liposomes containing PBS served as controls (7).

#### NETs quantification

MPO-DNA complexes as well as citrullinated Histone 4 were identified using capture ELISA as previously described (20, 29).

#### Histochemistry

Lungs were isolated and immediately fixed in 4% paraformaldehyde at 4°C for 6 hrs. For better cryopreservation, tissue was placed in 10%, 20% and 30% sucrose, tissue sinks embedded in optical cutting temperature (OCT) compound and stored at -80°C. Five µm thick lung slices were prepared and analyzed to assess inflammatory infiltration, tissue damage and fibrosis as previously described (7, 14).

#### Statistics

Results were reported as mean ± standard error of the mean (SEM), unless otherwise indicated. Kruskal-Wallis test followed by Dunn’s multiple comparison test was used to compare continuous variables between groups. All tests were two-sided and p values lower than 0.05 were considered statistically significant. In all analyses GraphPad Prism 10.3.1 was used

## Acknowledgments

The authors wish to thank MC Panzeri for electron microscopy, C. Covino for confocal microscopy and A. Fiocchi for histochemical analysis of the lung. Confocal and electron microscopies were carried out in the Advanced Light and Electron Microscopy BioImaging Center (ALEMBIC), an advanced microscopy laboratory established by the San Raffaele Scientific Institute and the Vita-Salute San Raffaele University. We thank the patients and the healthy donors who have provided the blood for this study.

## Funding

Funded by the European Union - Next Generation EU - NRRP M6C2 - Investment 2.1 Enhancement and strengthening of biomedical research in the NHS. Project PNRR-MR1-2022-12376638.

## Author contributions

AAM and NM defined the experimental strategy, performed experiments and wrote the manuscript. AC and AM performed experiments in the animal model. GAR and AAM selected the patients, elaborated the clinical database and performed statistical analysis. MEB, PRQ and MMC discussed the experimental strategy and corrected the manuscript.

## Competing interests

M.E.B. is the founder and part-owner of HMGBiotech (http://hmgbiotech.eu), a company that provides goods and services related to HMGB proteins. All other authors declare that they have no competing interests.

## Notes

### Competing Interest Statement

The authors have declared no competing interest.

